# Decoding transcriptional regulation via a human gene expression predictor

**DOI:** 10.1101/2020.04.07.029470

**Authors:** Yuzhou Wang, Yu Zhang, Jiazhen Gong, Jianqiang Bao, Shisong Ma

**Affiliations:** Hefei National Laboratory for Physical Sciences at the Microscale, School of Life Sciences, Division of Life Sciences and Medicine, University of Science and Technology of China, Innovative Academy of Seed Design, Chinese Academy of Sciences, Hefei, China; The First Affiliated Hospital of USTC, Division of Life Sciences and Medicine, University of Science and Technology of China, Hefei, China; School of Data Science, University of Science and Technology of China, Hefei, China

## Abstract

Transcription factors (TF) regulate cellular activities via controlling gene expression, but a predictive model describing how TFs quantitatively modulate human transcriptomes was lacking. We constructed a universal human gene expression predictor and utilized it to decode transcriptional regulation. Using 1613 TFs’ expression, the predictor reconstituted highly accurate transcriptomes for samples derived from a wide range of tissues and conditions. The predictor’s broad applicability indicated it had recapitulated the quantitative relationships between TFs and target genes ubiquitous across tissues. Significant interacting TF-target gene pairs were then extracted from the predictor and enabled downstream inference of TF regulators for diverse pathways involved in development, immunity, metabolism, and stress response. Thus, we present a novel approach to study human transcriptional regulation following the “understanding by modeling” principle.

## INTRODUCTION

The human tissues and cells are extremely complex entities, whose functional and physiological status are largely determined by transcriptomes. Understanding predictively and quantitatively how gene expression are regulated across diverse tissues and conditions is crucial to decode the genetic blueprint of human. As transcription is mainly modulated by transcription factors (TF), gene regulatory networks (GRN) that connect TFs and their target genes are important tools to decipher transcriptional regulation (*1–3*). To construct GRNs, TFs binding specificities to target genes have been determined systematically via charactering TF binding motifs and profiling genome-wide TF-DNA interactions (*4, 5*). GRNs were also deduced from transcriptome changes after perturbing TFs’ expression, using methods like Perturb-Seq (*6*). GRNs were further reverse-engineered from transcriptomes via computational algorithms like probabilistic graphical model, neural network, decision tress, LASSO regression, and network component analysis (*7–11*). However, to our knowledge no universal model has been built that can quantitatively predict human transcriptomes based on TF activities. Previously we developed a systematic approach named EXPLICIT (<u>E</u>xpression <u>P</u>rediction via <u>L</u>inear <u>C</u>ombination of <u>T</u>ranscription Factors) to construct a gene expression predictor for the model plant *Arabidopsis thaliana* (Geng et al., submitted). Here, we applied the same approach to build a universal human gene expression predictor. The predictor successfully reconstituted highly accurate transcriptomes for samples across a wide range of human tissues and cell types. It further enabled downstream inference of transcriptional regulators for human genes and pathways involved in development, immunity, metabolism, and stress response.

## RESULTS AND DISCUSSION

### A human gene expression predictor

Using the EXPLICIT approach (Geng et al., submitted), a human gene expression predictor was built to predict gene expression levels based on TFs’ expression. Large-scaled RNA-Seq-based human gene expression data were obtained from the ARCHS4 (v7) database (*12*), which contained transcriptomes for more than 167000 samples. After removing single-cell RNA-Seq (see below) and low-quality samples, 59097 samples were kept for the predictor model training. The predictor, constructed via ordinary linear least squares regression, shall use 1613 TF genes to predict the expression of 23346 non-TF genes (Figure 1A). Similar to the Arabidopsis predictor model, to obtain an accurate human predictor model, the number of samples within the training dataset was critical. Increasing the number of training samples effectively increased the model’s performance on predicting novel samples, as judging from the Pearson’s correlation coefficient (*r*) between the predicted and actual expression values ofthe independent test samples (Figure 1B). The over-fitting or optimism within the model also decreased with increasing training samples, as indicated by the decreased difference between the training and testing *r*. Thus, all 59097 human RNA-Seq samples were used to train the predictor model, which reduced over-fitting to a minimal level.

**Figure 1.**
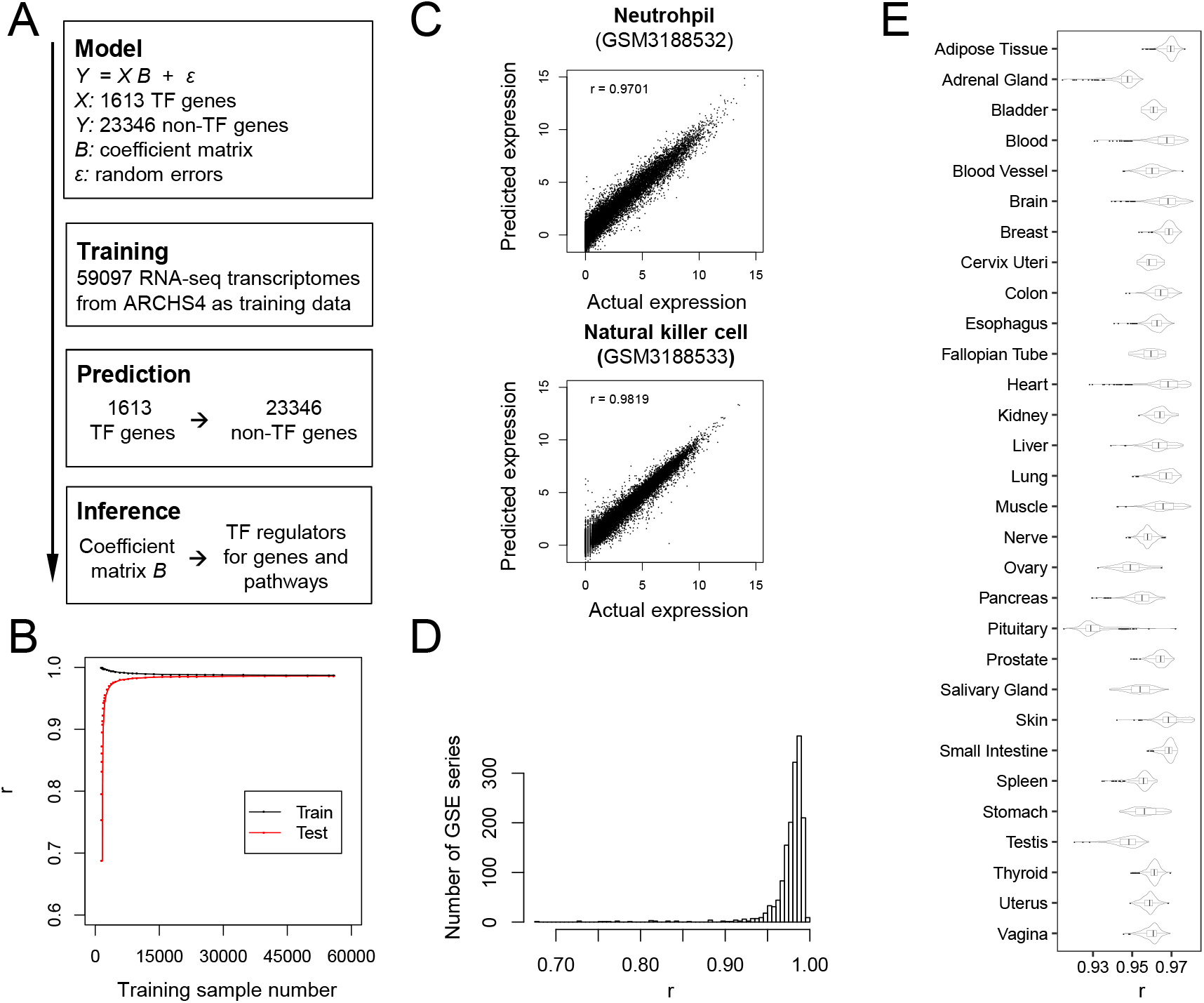
A human gene expression predictor and its predicting performance. **(A)** The analysis workflow. **(B)** Increasing training sample numbers improved the predictor’s performance. Predictor models trained with different numbers of training samples were tested for their predicting performance. Shown are the average Pearson’s correlation coefficients (*r*) between the predicted and actual transcriptomes for the training (black) and independent test samples (red, n=3000). **(C)** Leave-one-out cross-validation (LOOCV) test result for GSE115736 (*13*). The scatter plots show the log-transformed gene expression values for the predicted and actual transcriptomes of two samples. **(D)** LOOVC test results for 1545 GSE series contain 10+ samples. LOOCV test was conducted for each series, and the average *r* between predicted and actual transcriptomes for all its samples were calculated. The average *r* of every series were then tallied together to generate the histogram to show their distributions. **(E)** The predictor’s performance on predicting the GTEx transcriptomes. Boxplots and violin plots for the distribution of *r* between predicted and actual transcriptomes were shown for samples according to tissue types. 17382 samples in total, with between 9 to 2642 samples for each tissues.

The resulted human gene expression predictor was tested for predicting performance on novel samples via leave-one-out cross-validation (LOOCV). The training samples of the predictor originally came from 4000+ NCBI gene expression omnibus series (GSE), with each series being an independent study. In each LOOCV run, a single GSE series was selected and its samples were withheld, while the model was retrained with samples from all other series and then tested on the selected hold-out GSE series. The hold-out samples were deemed as novel independent samples for the retrained model. Shown in Figure 1C are the LOOCV test results for GSE115736 that had measured the transcriptomes of 12 different human hematopoietic cell types (*13*). Using 1613 TFs’ expression values, the predictor accurately predicted other 23346 non-TF genes’ expression in two samples of neutrophils (GSM3188532) and natural killer cells (GSM3188533), with *r* of 0.970 and 0.982 between the predicted and actual expression, respectively. The average *r* for all samples with the series was 0.980. LOOCV test was conducted for all 1545 GSE series that have 10+ samples. These series covered a broad range of experiment conditions used for human bulk RNA-Seq analysis, including sampling on different tissues or cell types. Despite such diversities, the predictor still achieved high accuracy, with an average *r* of 0.976±0.026 for all tested GSE series (Figure 1D).

The predictor was further tested on the GTEx (v8) dataset that have measured 17382 transcriptomes from diverse human tissues (*14*). The ARCHS4 training dataset did not contain samples from GTEx, and the two datasets were independent from each other. The predictor accurately predicted the GTEx transcriptomes derived from different tissues. Most tissues had average *r* ranging between 0.95 and 0.97, i.e 0.961 for thyroid, 0.964 for liver, and 0.969 for adipose tissue. Even the two tissues with the lowest accuracy, pituitary and adrenal gland, had average *r* of 0.931 and 0.946, respectively (Figure 1E). But the predictor had relatively low performance on single-cell RNA-Seq samples (data not shown), and these samples were removed before model training. Thus, the predictor model can be universally applied to most human bulk RNA-Seq samples from various tissues or cell types, indicating it had recapitulated the quantitative relationships between TFs and target genes ubiquitous across tissues. We envisioned such a model can be used to infer TF regulators for human genes and pathways.

### Human gene co-expression modules

To infer TF regulators for human pathways, we first identified gene co-expression modules containing genes that might function in the same pathways. A human gene co-expression network based on the graphical Gaussian model (GGM) was built using the ARCHS4 dataset via a previously published procedure with minor modification (*15*). After calculating the partial correlation coefficients (*pcor*) between all genes, 217686 co-expressed gene pairs with *pcor* >= 0.045 were selected to construct a gene co-expression network, which accounted for 0.07% of all possible gene pairs (Figure S1 and Table S1). The network, containing 24867 genes, was then assembled into 1416 gene co-expression modules (Figure 2A and Table S2). Via guilt-by-association, the genes within the same module co-expressed together and might have functions in the same pathways. Gene ontology (GO) analysis indicted 429 modules have enriched GO terms (*P* <= 1E-03) (Table S3).

**Figure 2.**
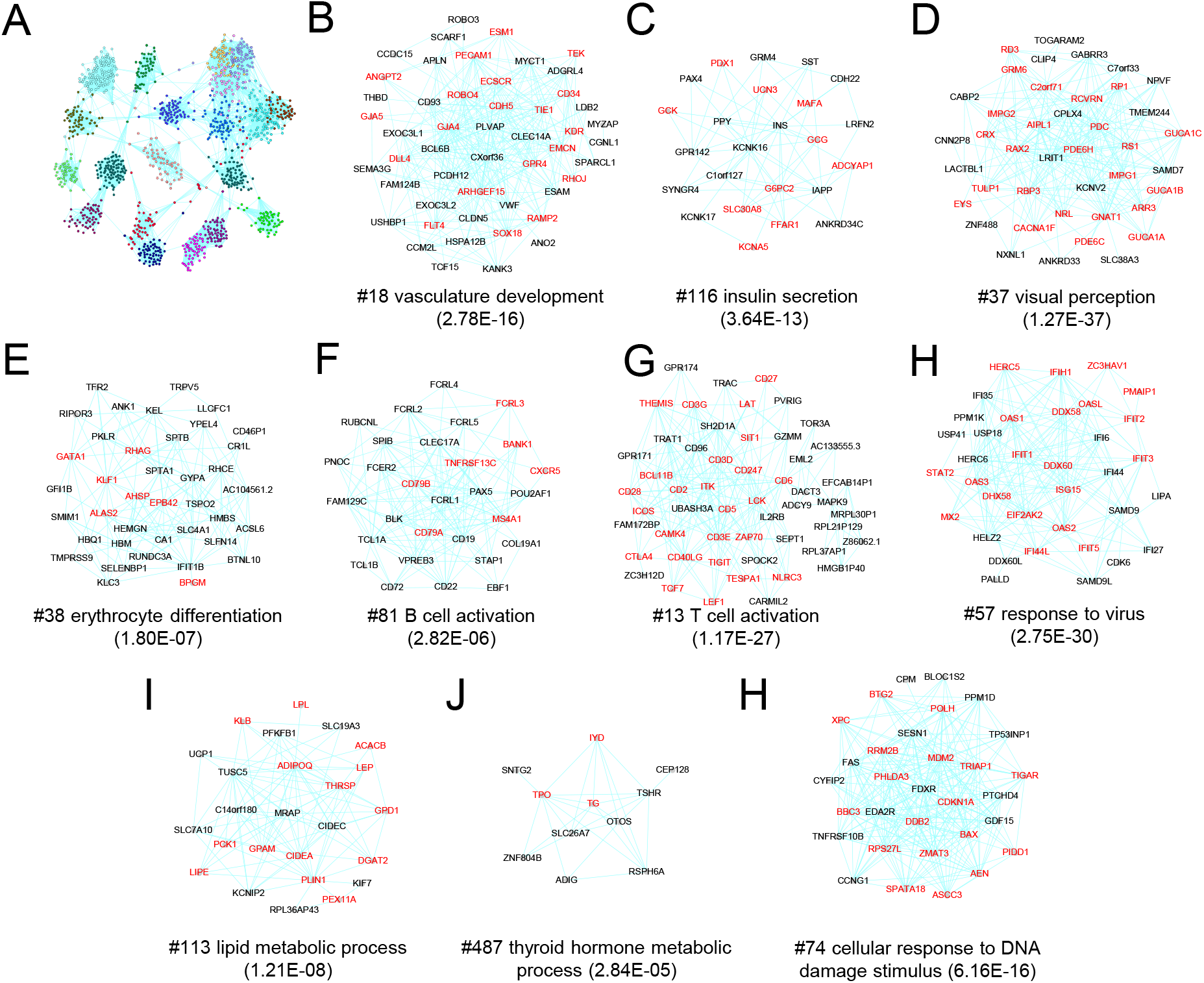
Human gene co-expression modules identified through co-expression network analysis. **(A)** The 20 largest module within the human GGM gene co-expression network. Dots represent genes and are color-coded by their module identities. Edges represent co-expression between genes. Due to space limit only the 20 largest modules are shown. **(B – K)** Subnetworks for selected gene co-expression modules. The module id number, its represented enrich GO term (and the enrichment p-value) are labeled for each module. Highlighted in red text are the genes possessing the represented GO term.

Further inspection pinpointed modules related to various development, immunity, metabolism, or stress response pathways. For example, a number of modules are involved in the development or functions of specific organs or tissues. Module #18 works in angiogenesis as it is enriched with 20 vasculature development genes (*P* = 2.78E-16), including *KDR* and *TEK* (*16*), #116 functions in insulin secretion (*P* = 3.64E-13) with biased expression in pancreas, while #37 is involved in visual perception (*P* = 1.27E-37) and photoreceptor cell development (Figure 2B-D, and S2). Modules related to hematopoietic and immunity cells were also discovered, including #38 for erythrocyte differentiation, #81 for B cell activation, #13 for T cell activation, and #57 for defense response to virus via type I interferon signaling pathway (Figure 2E-H). Also discovered were modules functioning in metabolism, such as #113 for lipid metabolism with biased expression in adipose tissues and #487 for thyroid hormone metabolism (Figure 2J, K, and S2). Stress response modules were also revealed, such as #74 involved in DNA damage response and p53 mediated apoptotic signaling pathway (Figure 2L). Thus, the network analysis identified gene modules for a broad range of pathways. As genes within the same module have similar co-expression patterns, they might be regulated by the same transcription factors. Identifying such TFs will be highly valuable since these modules play critical roles in diverse developmental and physiological processes related to illnesses like obesity, diabetes, tumors, virus infection, and cardiovascular, immunity, or eye diseases.

### TF regulators for genes and pathways

The human gene expression predictor was then used to infer TF regulators for genes and pathways, following the EXPLICIT approach (Geng et al., submitted). Individual regression coefficients within the predictor’s coefficient matrix *B* were tested for significance, and 2286331 coefficients with p-values <= 1E-12 (approximately equal to a Bonferroni-adjusted *P* of 5E-05) were deemed as significant and their corresponding TF-gene pairs extracted (Table S4). These significant interacting TF-gene pairs connected TFs to their target genes, and the TFs connected with a target gene were deemed as that gene’s predictor TFs. In average each gene has 98 predictor TFs, which could also be considered as the gene’s potential transcriptional regulators. For example, the gene *GNAT1*, encoding the rod-specific transducin α subunit functioning in phototransduction (*17*), had 101 identified predictor TFs (Figure 3A). Among the top predictor TFs were three known regulators of *GNAT1* expression, *NRL, NR2E3*, and *CRX*, while the rest might also contain additional regulators as many of them regulate retinal development (*18*).

TF regulators were also inferred for gene co-expression modules at the pathways level. For instance, *GNAT1* is contained within Module #37 that function in visual perception. The module has 36 non-TF genes in total. Predictor TFs for the module were identified as the common predictor TFs shared by these genes. Specifically, if a TF was a common predictor TF for at least 6 as well as 20% of the non-TF genes within a module and its target genes were over-represented within the same module (*P* <= 1E-06), the TF was considered as a predictor TF for that module. *CRX* is a predictor TF for 32 genes within Module #37, while it has only 712 target genes among all 23346 non-TF genes, thus its target genes are enriched 29 folds within the module (*P* = 8.3E-45). Therefore, *CRX* was considered as a predictor TF for Module #37. In total 23 predictor TFs were identified for the module (Figure 3B and Table S5). Surprisingly, 16 of them have been shown before to be regulators of retinal development or functions (Table S6, same as below for other modules), such as *CRX, NRL, NR2E3*, and *OTX2 (19*). Similarly, 24 predictor TFs were identified for Module #18 involved in vasculature development, among which are 15 known regulators of angiogenesis, like *SOX17* and *SOX18* (Figure 3C) (*20*). And for the insulin secretion and pancreas related Module #116, 31 predictor TFs were identified, 19 of which are known regulators of pancreas development or functions, i.e. *PDX1, PAX6, MAFA* and *NEUROG3* (Figure 3D) (*21*). Thus, among the identified predictor TFs, more than 60% are known regulators, while the rest might be potential novel regulators.

**Figure 3.**
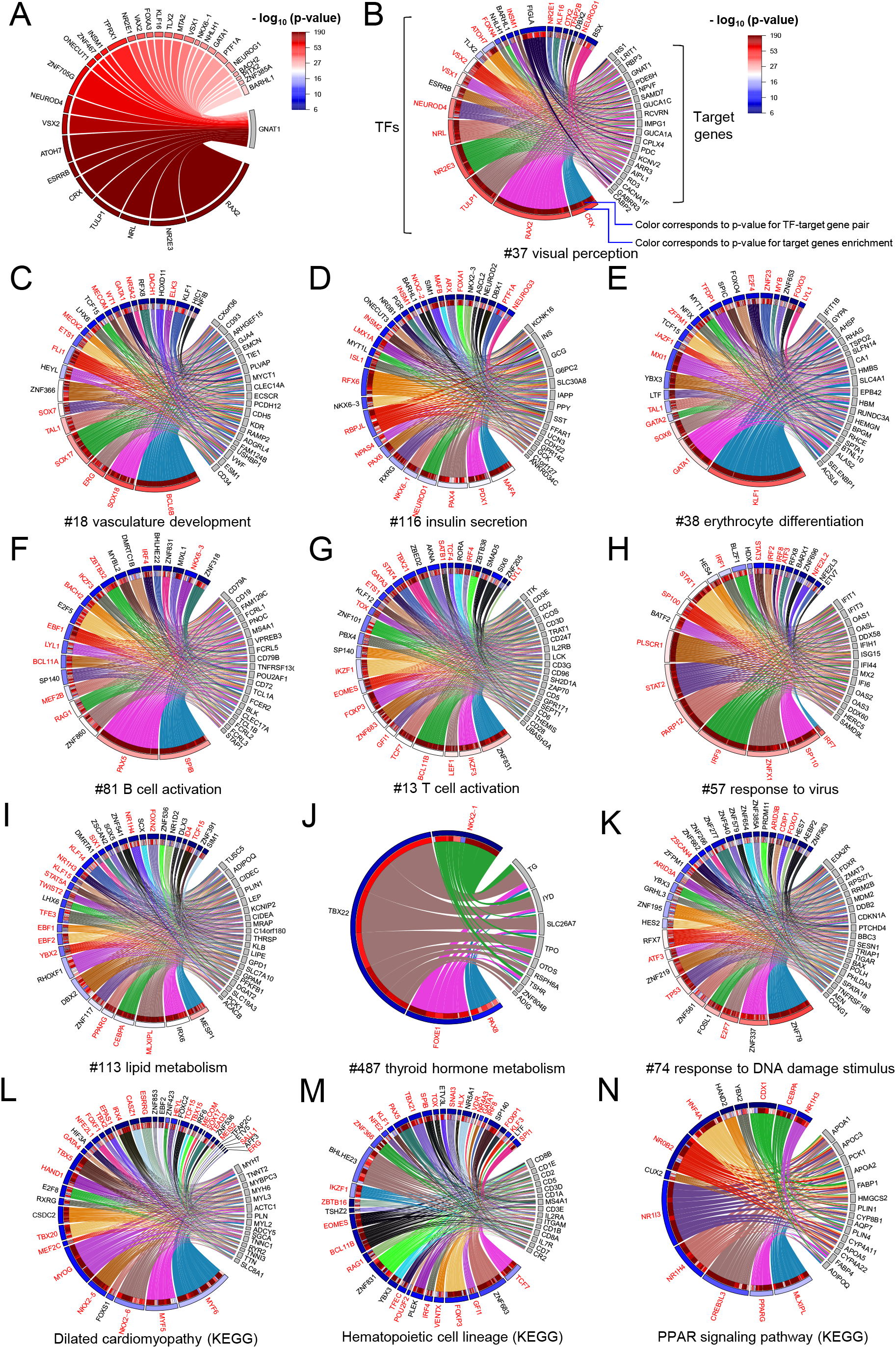
Predictor TFs for genes and pathways identified from the predictor. **(A)** Interactions between *GNAT1* and its predictor TFs are shown as a Chord diagram. Colors indicate the p-values for the coefficients corresponding to the TF-target gene pair, and the links’ widths are proportional to the coefficients’ magnitude. Only top 30 predictor TFs are shown due to space limit. **(B-H)** Chord diagrams connecting predictor TFs to their target genes within the selected gene coexpression modules. Links are colored by predictor TF genes, their left ends’ colors indicates p-values of the corresponding coefficients, while their widths are proportional to the coefficients’ magnitude. Enrichment of the predictor TFs target genes within the module are also indicated by colors. TFs highlighted in red text are known TF regulators. Only 15-20 target genes were shown due to space limit. **(L-N)** Chord diagrams connecting predictor TFs to their target genes within KEGG pathways.

Predictor TFs were also recovered for the modules involved in erythrocyte differentiation (#38), B-cell activation (#81), T-cell activation (#13), and defense response to virus (#57) (Figure 3E-H). Module #38 has 22 predictor TFs, 14 of which are known regulators of erythrocyte differentiation or functions, like *KLF1, GATA1* and *TAL1 (22*). Module #81 has 21 predictor TFs, 12 of which have known functions in B-cells, i.e. *PAX5, EBF1* and *BCL11A* (*23*). Module #13 has 30 predictor TFs, and 18 of them are known regulators of T cell development or functions, such as *TCF7, GATA3*, and *BCL11B (24*). Similarly, 15 out of 24 predictor TFs for Module 57 are also known regulators of defense response to virus, such as *STAT1, STAT2, IRF7* and *IRF9* (*25*). Again, more than 55% of the identified predictor TFs are known regulators of the pathways involved.

Also identified were predictor TFs for the modules participating in lipid metabolism (#113), thyroid hormone metabolism (#487), and DNA damage response (#74) (Figure 3I-K). Module #113 has 33 predictors, 17 of them are known regulators of lipid metabolism or adipogenesis, including *PPARG* and *CEBPA* that modulate adipocyte differentiation (*26*). Similarly, Module #487 has 4 predictor TFs, 3 of which (*PAX8, FOXE1*, and *NKX2-1*) are among the four curial TFs regulating thyroid development and their mutations are associated with congenital hypothyroidism (*27*). Noted that the other crucial TF *HHEX* also had 6 target genes within the module, but its enrichment p-value (5.98E-05) was slightly larger than the selected cutoff. In contrast, Module #74 has 30 predictor TFs, but only 8 of them are known regulators of DNA damage response, including the master regulator *TP53* (*28*). Interestingly, the module’s predictor TFs are over-represented with 15 zinc finger protein genes, but only *ZSCAN4* has known functions in DNA damage response (*29*), while the rest genes’ relevance remains to be investigated, such as the top predictor TFs *ZNF79, ZNF337*, and *ZNF561*.

Including the ones mentioned above, predictor TFs were identified for 757 co-expression modules, with each module having 8.6 predictor TFs in average (Table S5). Besides co-expression modules, predictor TFs were also identified for annotated KEGG pathways (Table S7) (*30*). For example, the dilated cardiomyopathy (DCM) pathway (hsa05414) contains 96 human genes related to DCM, 94 of which are among the non-TF genes used for our model training. Treating these 94 genes as a group, 40 predictor TFs were identified, among which 27 are known regulators of heart or muscle development, such as *NKX2-5, GATA4, HAND1*, and *TBX5* (Figure 3L) (*31*). Similarly, predictor TFs were identified for the hematopoietic cell lineage (hsa04640) and PPAR signaling (hsa04979) pathways, with >= 65% of them being known regulators (Figure 3M, N). These two pathways play critical physiological roles by regulating blood cells regeneration and modulating lipid and glucose metabolism and overall energy homeostasis, respectively (*32, 33*).

### Verification of predictor TFs by independent gene expression dataset

The above analysis identified a number of predictor TFs as novel potential regulators for different pathways whose functions remained to be investigated. However, a question remains that if finding these predictor TFs depended on a specific training dataset, such as ARCHS4. To address that, an independent human gene expression predictor was also built using the GTEx (v8) dataset. As GTEx had much less samples than ARCHS4, far less significant TF-target gene pairs were recovered. To make the two models more comparable, significant TF-target gene pairs were extracted from the coefficients with p-value <= 1E-09 (approximately equal to a Bonferroni-adjusted *P* of 0.05) from the GTEx-based predictor. In average, only 13.3 predictor-TFs were identified for each target gene. Nevertheless, using the GTEx-based predictor, similar predictor TFs were still identified, although their numbers were much less (Table S8). For example, 7 predictor TFs were identified for Module #74 of DNA damage response, including 3 known regulators (*TP53, CDIP1*, and *E2F7*) as well as 4 uncharacterized TFs (*ZNF79, ZNF561, RFX7*, and *ZNF337*) (Figure 4A). Interestingly, these 4 uncharacterized TFs, including 3 zinc finger protein genes, were also identified via the ARCHS4-based predictor. Thus, these predictor TFs were identified twice by two independent datasets. Similar uncharacterized predictor TFs verified by two independent datasets were also recovered for other modules involved in lipid metabolism, erythrocyte differentiation, T cell activation, response to virus, and vasculature development (Figure 4B-E). These predictor TFs can be treated as prioritized candidate genes for future potential regulator research. Additionally, it should be noted that the GTEx-based predictor has much less training samples and statistics power than the ARCHS4-based predictor, which might limit its ability to verify more uncharacterized predictor TFs.

**Figure 4.**
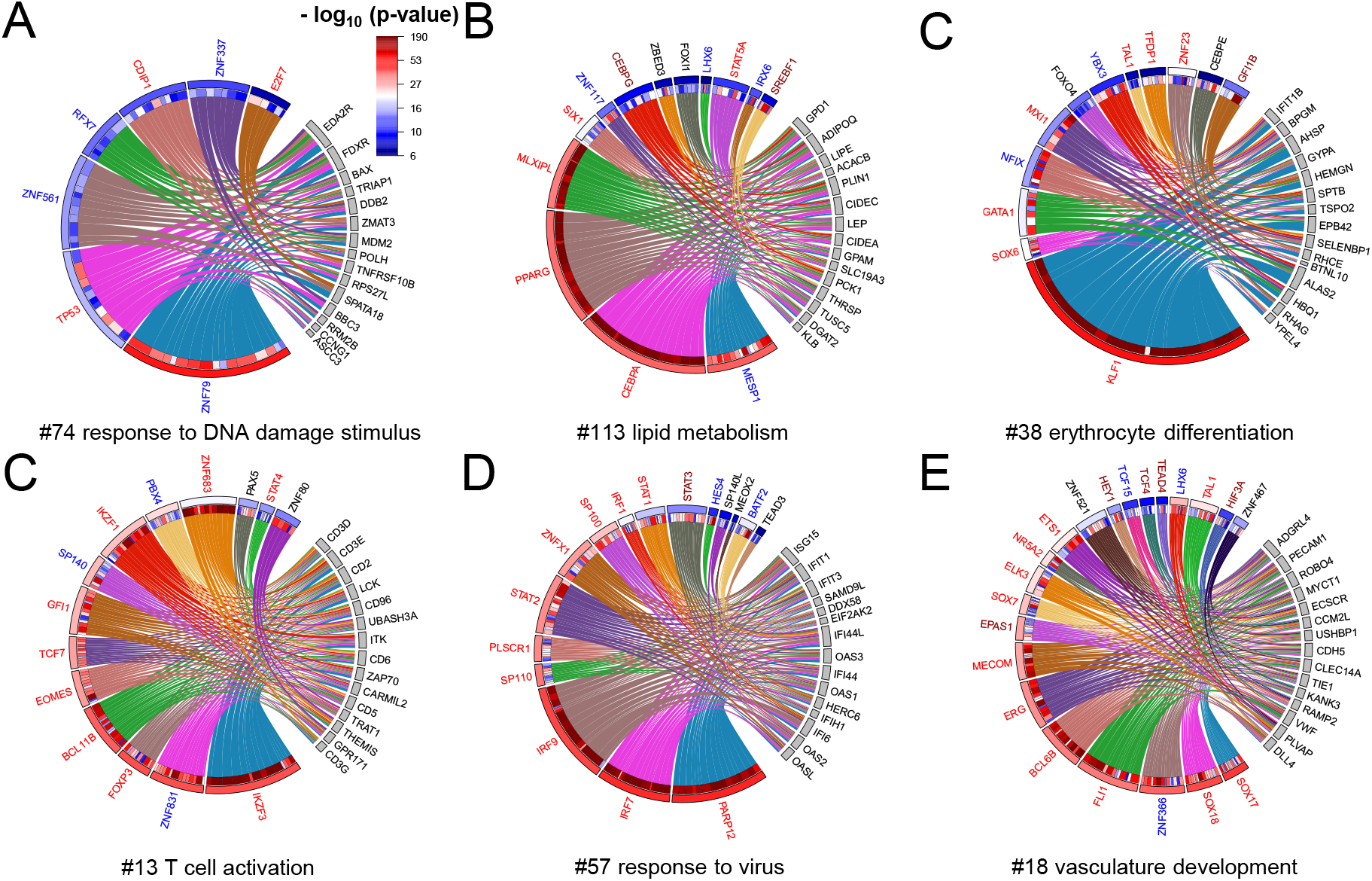
Predictor TFs for selected modules verified by an independent predictor trained with the GTEx v8 dataset. Predictor TFs for modules were identified via the GTEx-based gene expression predictor, as shown in the Chord diagrams (A-E). Legends are the same as Figure 3, except that TFs in blue text are uncharacterized TFs identified by both predictors, TFs in red text are known TF regulators identified by both predictors, and TFs in dark-red text are known TFs identified by the GTEx-based predictor only.

Thus, by using large-scaled transcriptome dataset, we were able to construct a universal human gene expression predictor. There were reports before on predicting transcriptomes using a selected number (100 - 300) of informative genes, which served as the basis to develop cheaper alternatives rather than RNA-Seq to measure gene expression profiles (*34, 35*). Our analysis had different goals. By using almost all TFs as regressors to construct the predictor model, our predictor allowed downstream inference of TF regulators in addition to predicting transcriptomes. The analysis followed the “understanding by modeling” principle, which is similar to the methodology of “understanding by building” in Synthetic Biology (*36*). By building a universal gene expression predictor model, it became possible to reconstitute highly accurate transcriptomes using only TF genes’ expression for samples derived from a wide range of tissues or cell lines. The broad applicability and high accuracy of our predictor model indicated there exist a set of ubiquitous and tissue-independent quantitative relationships between TFs and target genes, which had been recapitulated and abstracted by the predictor. These quantitative relationships can be translated into regulatory interactions in many cases and revealed transcriptional regulators for genes and pathways, as demonstrated by the aforementioned examples. Thanks to its tissue-independence, the predictor enabled the identification of TF regulators for completely different tissues or cell types, i.e. heart, pancreas, thyroid, adipose, and different hematopoietic cell lines. Besides gene coexpression modules, the predictor also inferred regulators for annotated KEGG pathways and possibly for any pre-defined gene sets, including those related to human diseases. The predictor is expected to facilitate mechanistic studies on human transcriptional regulation.

## MATERIALS AND METHODS

### Gene expression data and gene expression predictor

A large-scaled processed RNA-Seq-based human transcriptome dataset was downloaded from the ARCHS4 (v7) database (https://amp.pharm.mssm.edu/archs4/data.html) (*12*). The downloaded file (human_transcript_v7.h5) contained read counts for 178136 transcripts in 167726 RNA-Seq samples, which originated from 6000+ NCBI GSE series. The GSE series containing single-cell RNA-Seq samples were manually removed. The transcript counts were then consolidated according to gene identities to produce a gene counts table, from which cpm (counts per million) gene expression values were calculated. The samples with less than 1000000 mapped reads or with reads mapping rates < 60% or with less than 5000 genes having expression values cpm >= 1 were removed. Further filtered out were the GSE series with > 50% samples removed in previous step. There were a limited number of transcriptomes generated using tissue samples from the GTEx project, which were also removed from the ARCHS4 training dataset. Finally, 59097 samples from 4214 GSE series were remained. Among these remaining samples, the low-expressed genes that have cpm values >= 1 in less than 100 samples were removed. It was found out that 1998 genes have duplicated expression values across samples with other genes, and they were also removed. In order to use the GTEx (v8) dataset, which was processed via a different pipeline (*14*), to verify the predictor, only the genes common to both datasets were kept. As a result, 24959 genes were remained. The cpm gene expression matrix for these 24959 genes in 59097 selected samples were then log-transformed via log2(cpm+1) and kept for further analysis.

Two gene expression matrices, *X* for 1613 TF-genes and *Y* for 23346 non-TF genes, were then extracted according to a human TF gene list obtained from AnimalTFDB 3.0 (*37*). They were used to build a human gene expression predictor via the EXPLICIT approach described before (Geng et al, submitted). Briefly, a linear regression model, *Y* = *X B* + *ε*, was trained via the least squares method. The significances of individual coefficients within the predictor model were determined via hypothesis testing method for multiple linear regression (*38*). Significant interacting TF-target gene pairs were extracted corresponding to the coefficients with p-value less or equal to selected cutoff and used to identify predictor TFs for non-TF genes as well as gene modules.

To test the model’s performance, two actual expression matrices, *X_t_* for TFs and *Y_t_* for non-TFs genes, were extracted from the test samples’ actual transcriptomes, and a predicted expression matrix *Y_p_* was estimated as *Y_p_ = X_t_ B*. Note that for LOOCV test, the predictor model was retained with all other samples without the test samples. *Y_t_* and *Y_p_* were then used to compute Pearson’s correlation coefficients between actual and predicted transcriptomes for the individual samples they contained. A MATLAB script developed in-house was used to conduct the analysis.

As another independent dataset, processed and publically available RNA-Seq-based human transcriptome data were also obtained from the GTEx (v8) project (https://gtexportal.org) (*14*). The downloaded file (GTEx_Analysis_2017-06-05_v8_RNASeQCv1.1.9_gene_reads.gct.gz) contained read counts for 56200 genes in 17382 samples spanning 54 human tissue-sites. The gene counts expression table was used to calculate the cpm gene expression values, which were subjected to log-transformation via log_2_(cpm+1). Two expression matrices, containing the same TF and non-TF genes in the same order as the ARCHS4 training dataset respectively, were then extracted and used for predictor performance testing. These two expression matrices were further used to train an independent GTEx-based human gene expression predictor, so as to compare with the ARCHS4-based predictor to investigate if they recover similar predictor TFs for gene co-expression modules.

### Gene co-expression modules and predictor TFs identification

Using the ARCHS4 gene expression matrix containing 24959 genes in 59097 samples, a human GGM gene co-expression network was constructed via a previously published procedure with minor modification (*15*). Briefly, *pcors* between all gene pairs were calculated via a random sampling procedure that consisted of 20000 iterations. In each iteration, 2000 genes were randomly selected and used for partial correlation coefficient (*pcor*) calculation. Previously, a shrinkage approach implemented in the GeneNet package in R was employed to estimate the covariance matrix between the selected genes and use it for *pcor* calculation (*39*). The shrinkage approach was developed specifically to address the “small n, large p” problem common to biological transcriptome datasets, which usually had far less samples number (n) than genes number (p). In the current study, because the samples number (59097) is much larger than the selected genes number (2000), the sample covariance estimator were directly used to estimate the covariance matrix for the selected genes. The *pcors* between the selected genes were then calculated, which was related to inverse of the covariance matrix (*39*). After 20000 iterations, each gene pairs were randomly sampled together in 128 iterations with 128 *pcors* calculated, and the *pcor* with the lowest absolute value was then chosen as that gene pair’s final *pcor*. The gene pairs whose *pcors* were >= 0.045 were then chosen to construct a GGM gene co-expression network for human. A MATLAB script was developed in-house to conduct the analysis. The network was then clustered into 1416 gene modules that contain at least 6 genes via the MCL algorithm. The network and its modules were visualized in Cytoscape (V3.4.0) (*40*). Human GO annotations were obtained from Ensembl BioMart (http://www.ensembl.org/biomart/martview) and used for GO enrichment analysis via the hypergeometric distribution.

Predictor TFs were then identified for the modules, following EXPLICIT approach used for the Arabidopsis study (Geng et al., submitted). Briefly, predictor TFs were first identified for individual genes from the significant interacting TF-target gene pairs out of the gene expression predictor model. For any gene module, if a TF was a common predictor TF for at least 6 as well as 20% of the non-TF genes within the module and its target genes were over-represented within the same module (*P* <= 1E-06) as determined via the hypergeometric distribution, the TF was considered as a predictor TF for that module. A perl script developed in-house was used to conduct the analysis. Chord diagrams were drawn using the ‘circlize’ package in R to illustrate the interactions between predictor TFs and target genes within a gene module (*41*). Besides gene coexpression modules, the gene sets for annotated pathways were also obtained from the KEGG database and used for predictor TFs inference (*30*).

## ACKNOWLEDGEMENT

This work was supported by grants from the National Natural Science Foundation of China (31770268), the Strategic Priority Research Program of the Chinese Academy of Sciences (XDA24010302), the Fundamental Research Funds for the Central Universities (WK2070000091), and University of Science and Technology of China (Start-up fund to S.M.). The numerical calculations in this manuscript were conducted on the supercomputing systems in USTC Supercomputing Center and USTC School of Life Sciences Bioinformatics Center.

## SUPPLEMENTAL FIGURES AND TABLES

**Figure S1.** A histogram showing the distribution of all *pcors* between 24959 genes. Only those gene pairs with *pcors* >= 0.045 were used for GGM gene co-expression network construction.

**Figure S2.** A heatmap showing the tissue specific expression patterns form the genes in Module #113 and #116. Shown are the median gene expression values for each tissue were calculated from the GTEx (v8) transcriptomes.

**Table S1.** 217686 significant co-expressed gene pairs used to construct the GGM gene coexpression network.

**Table S2.** Genes and their module identities within the co-expression network.

**Table S3.** GO enrichment analysis results for the identified gene co-expression modules.

**Table S4.** Significant interacting TF-target gene pairs identified form the human gene expression predictor.

**Table S5.** Predictor TFs identified for the gene co-expression modules.

**Table S6.** Known regulators among the predictor TFs identified for the selected gene co-expression modules.

**Table S7.** Predictor TFs identified for KEGG pathways.

**Table S8.** Predictor TFs identified for gene co-expression modules based on the GTEx-based predictor.

