## Supplementary figures and images for "Decoding transcriptional regulation via a human gene expression predictor"

### Figure S1

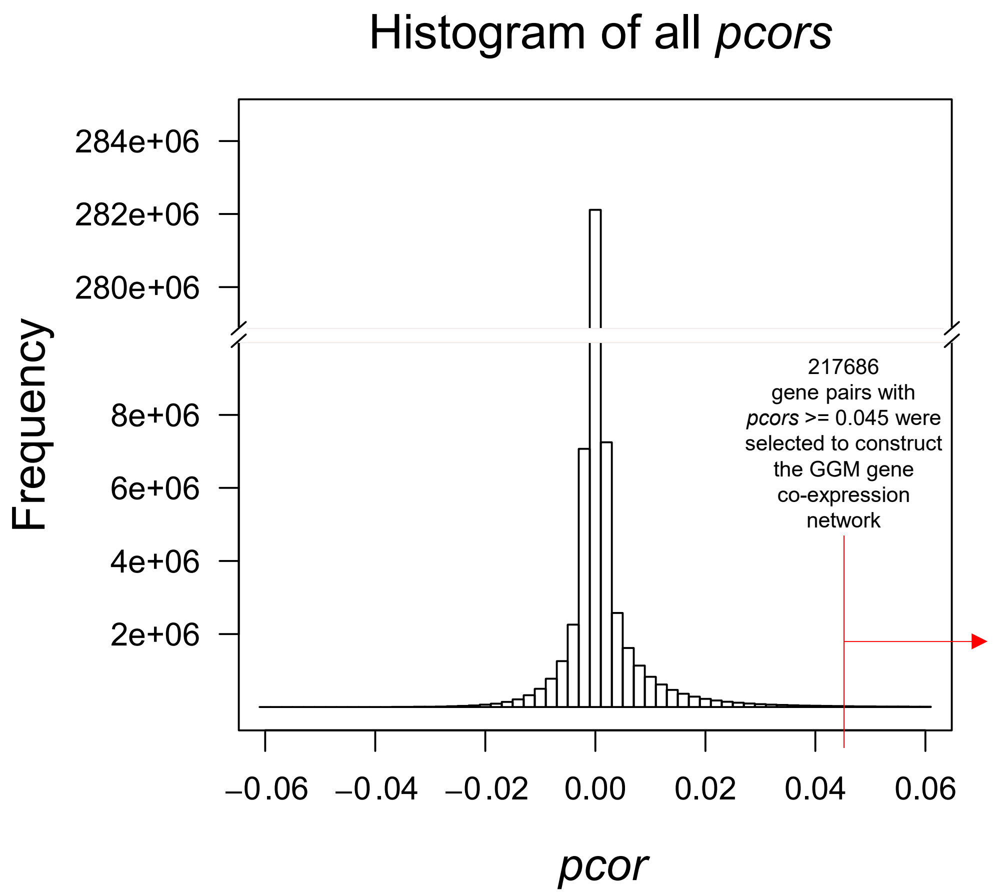

### Figure S2

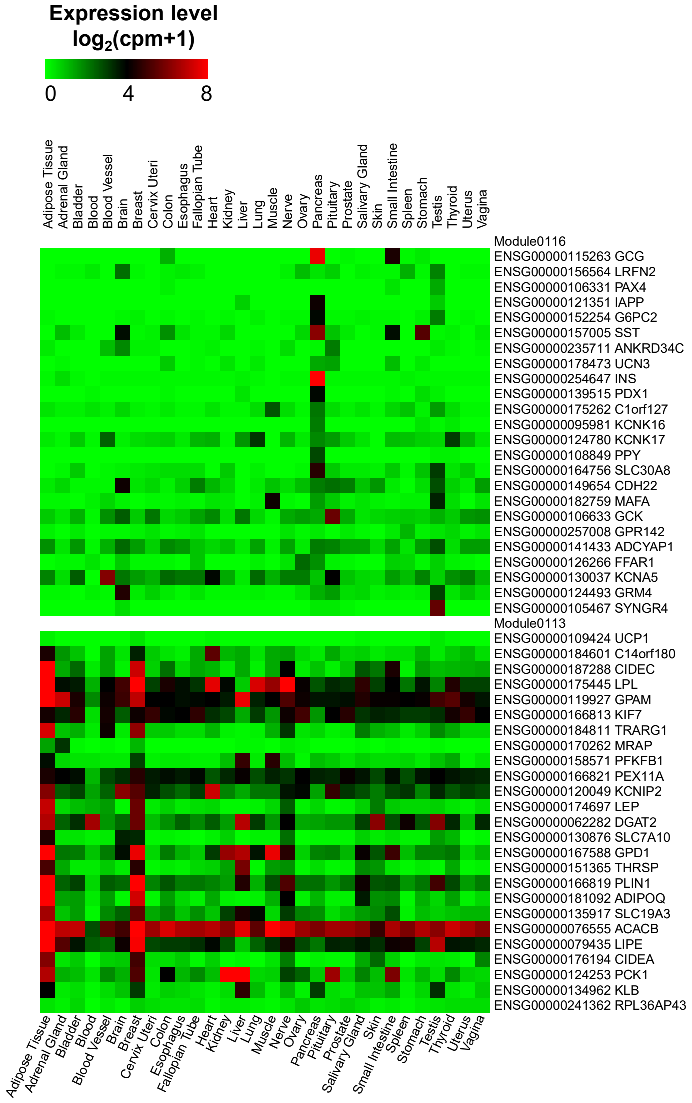
